# Succinylated lysine residue prediction revisited

**DOI:** 10.1101/2022.05.20.492505

**Authors:** Shehab Sarar Ahmed, Zaara Tasnim Rifat, Mohammad Saifur Rahman, M. Sohel Rahman

## Abstract

**Motivation:** Lysine succinylation is a kind of post-translational modification (PTM) which plays a crucial role in regulating the cellular processes. Aberrant succinylation may cause inflammation, cancers, metabolism diseases and nervous system diseases. The experimental methods to detect succinylation sites are time-consuming and costly. This thus calls for computational models with high efficacy and attention has been given in the literature for developing such models, albeit with only moderate success in the context of different evaluation metrics. One important aspect in this context is the biochemical and physicochemical properties of amino acids, which appear to be useful as features for such computational predictors. However, some of the existing computational models did not use the biochemical and physicochemical properties of amino acids, while some others used them without considering the inter-dependency among the properties.

**Results:** The combinations of biochemical and physicochemical properties derived through our optimization process achieve better results than the results achieved by the combination of all the properties. We propose three deep learning architectures, CNN+Bi-LSTM (CBL), Bi-LSTM+CNN (BLC) and their combination (CBL_BLC). We find that CBL_BLC is outperforming the other two. Ensembling of different models successfully improves the results. Notably, tuning the threshold of the ensemble classifiers further improves the results. Upon comparing our work with other existing works on two datasets, we find that we successfully achieve better sensitivity and specificity through varying the threshold value.

**Availability:** https://github.com/Dariwala/Succinylation-with-biophysico-and-deep-learning

## 1 Introduction

Lysine succinylation (addition of succinyl group to the lysine residue) is an important Post-translational modification (PTM) in regulating the cellular processes [7]. It can encourace change of charge in the surroundings and stimulate structural and functional adjustments to substrate proteins. Succinylation is involved in a variety of core energy metabolism pathways, including amino acid degradation, tricarboxylic acid cycle and fatty acid metabolism, TCA cycle, pentose phosphate pathway, glycolysis/gluconeogenesis, and pyruvate metabolism [20]. Hence, detecting and investigating the succinylated lysine residues (will be referred to as SLR henceforth) is the key to understand the function of proteins. Moreover, succinylation may lead to inflammation, tuberculosis [21]. The dysregulation of succinylations have also been found to cause diseases including cancers [23] and metabolism diseases [27]. Therefore, it is crucial to identify SLRs in the field of physiology too.

Although there exist several experimental techniques (e.g., mass spectrometry, liquid chromatography) for detecting PTM sites, they are usually costly and time consuming. This thus calls for in silico computational techniques to provide predictions of SLRs with high efficacy.

The first SLR predictor, named iSuc-PseAAC, was presented in [26]. It achieved a sensitivity of only 50%. Around the same time, another group independently developed another predictor named SuccFind [24]. They ranked the biochemical and physicochemical (will be referred to as biophysico henceforth) properties from AAindex [13] and adopted the best property, i.e., point. Another predictor, SucPred [30] also exploits sequence based features like SuccFind. They reported only sensitivity score of their proposed model.

One year later, iSuc-PseOpt [11] was proposed which adopted pseudo amino-acid composition (PseAAC) to encode the peptides surrounding the lysine residues. However, this work suffers from data leakage problem (i.e., information of the test set was used to preprocess the dataset). Within a very short period, the same authors came up with another predictor, pSuc-Lys [12], which was free from the data leakage issue.

In 2017 and 2018, Dehzangi *et al*. proposed four succinylation predictors SucStruct [14], PSSM-Suc [3], Success [15] and SSEvol-Suc [4]. All four classifiers made the same mistake done by the authors of iSuc-PseOpt [11].

On the other hand, Hasan *et al*. proposed three predictors in three consecutive years, namely, SuccinSite [7], SuccinSite2.0 [5] and GPSuc [6]. These three works applied several feature selection strategies to reduce the dimesionality of the feature space. However, the train dataset is not the same as the ones used by the previous works. So, the comparison made in these works with the existing works is not valid.

Psucce [16] used one-hot encoding and AAC in addition to top ten biophysico properties. Inspector [31] incorporated several sequence based features and achieved an improved sensitivity with respect to SuccinSite2.0, Psucce and GPSuc. A very recently developed tool, predML-Site [1] used several sequence-based features including autocorrelation function (ACF), several biophysico properties, pseudo amino acid composition (PseAAC) and used the SVM algorithm for training.

A novel deep-learning based predictor named MUscADEL [2] was developed using recurrent neural network (RNN) for predicting SLRs. But, nothing has been clearly mentioned about the independent dataset that was used to evaluate the performance and compare with other existing works.

Later on, HybridSucc [17] was developed by integrating traditional ML algorithms with Deep Neural Networks (DNNs). It achieved significantly better AUC values compared to several existing predictors.

One of the pioneering works in this field, DeepSuccinylSite [22], experimented with both one hot vectors and embedding vectors and fed them into a convolutional neural network This tool reported significantly better performance compared to previously mentioned predictors.

In 2021, LSTMCNNsucc [10] was proposed. It achieved a higher Matthews Correlation Coefficient (MCC) compared to DeepSuccinylSite. However, the achieved sensitivity is very low.

Finally, during the write-up stage of this thesis, the publication of a new predictor, called DeepSucc [29] came to our knowledge. While the reported performance is significantly better than all other previous predictors, upon careful scrutiny we found that the codebase shared by the authors contains multiple discrepancies. This will be elaborately discussed in Section 3.6.

## 2 Methods

### 2.1 Datasets

We use two datasets as follows (Table 1). The first dataset (D1) is collected from the Protein Lysine Modification Database (PLMD) [25]. The authors of [7] compiled the second dataset (D2) from UniprotKB/SwissProt and NCBI Protein Sequence Database. The detailed steps of preprocessing the datasets are described in Section 1 of the Supplementary File.

**Table 1.** Summary statistics of the two datasets, D1 and D2.

| Type | D1 |  | D2 |  |
| --- | --- | --- | --- | --- |
|  | Positive Samples | Negative Samples | Positive Samples | Negative Samples |
| Training | 5816 | 5826 | 3790 | 7460 |
| Validation | 696 | 686 | 960 | 2040 |
| Test | 1479 | 16457 | 254 | 2977 |

### 2.2 Representation of samples

We refer to the SLRs as positive samples and all other lysine residues in the same proteins as negative samples. Each of the lysine residue is represented by a number amino acids (say, *w*) on both its upstream and downstream (Figure 1 of Supplementary File). We will refer to 2*w* + 1 as the *context window*. We use 2*w* + 1 = 33 as has been used by DeepSuccinylSite [22]. If there are less number of amino acids on any side of the concerned lysine residue, mirror effect is used to keep the *context window* fixed for each sample.

**Fig. 1.**
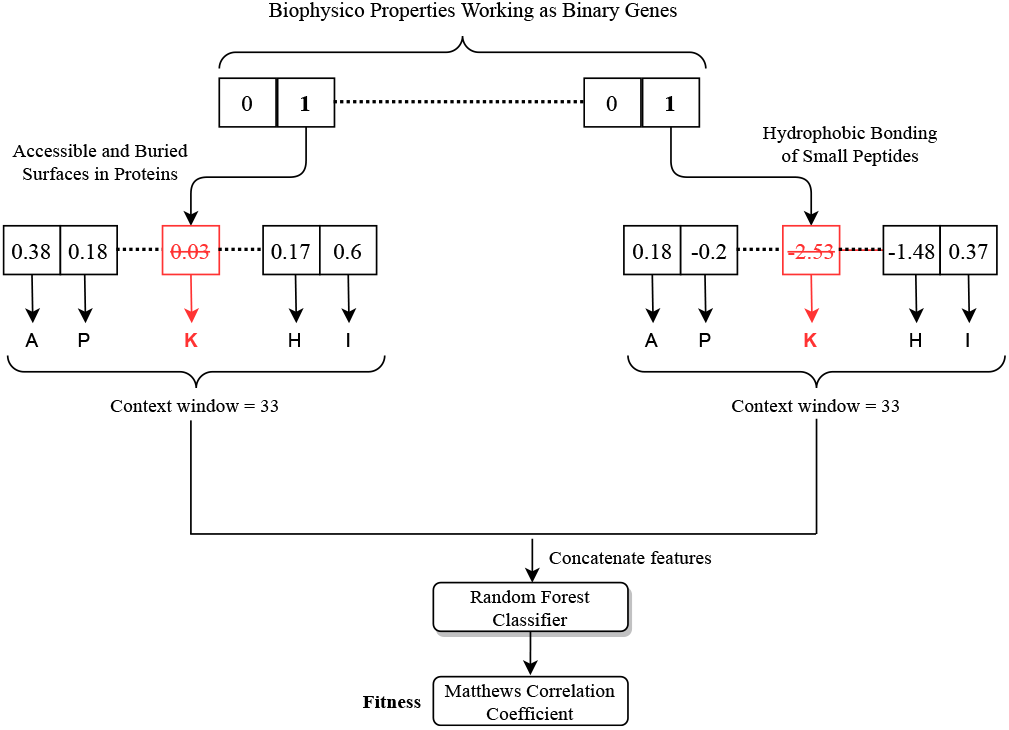
Evaluation of fitness value for an individual from the chromosome. An individual consists of a number of amino acids, the amino acids from the upstream and downstream of the lysine residue. The lysine residue itself is ignored because it is the same across all the samples. The biophysico properties corresponding to the binary genes having value 1 are considered for each of these amino acids. These features are concatenated and an RF is trained on the feature space of training samples. The MCC on the corresponding validation set is considered the fitness of the individual.

### 2.3 Selecting a subset of biophysico properties

While we are interested to include biophysico properties as features in our model, many of the 566 biophysico properties may not be relevant for the prediction of SLRs. Therefore, for each of them, a Random Forest (RF) classifier is trained with the training sets of both D1 & D2 by replacing each amino acid with its value for the corresponding property. The classifier is then evaluated on the corresponding validation dataset with respect to Matthews Correlation Coefficient (MCC) (See details about the MCC and other performance metrics in Section 2 of Supplementary File). The 95% percentile of the 566 MCC values is computed and only those features are considered for which the MCC values are greater than the 95% percentile. So, we are remained with 566 × 0.95 *≈* 29 properties for each dataset. In what follows, we only focus on these 29 properties.

### 2.4 Inheritable bi-objective genetic algorithm

Exhaustive search for the best combination is infeasible because there are 2^29^ *−* 1 possible non-empty combinations from the 29 biophysico properties. Therefore, we need to leverage heuristic method to search for suitable combinations in this big search space. Inheritable bi-objective genetic algorithm (IBCGA) [8] consists of an intelligent genetic algorithm [9] with an inheritable mechanism. The algorithm adopts a divide and conquer strategy with orthogonal array crossover (please refer to Section 3.2 of Supplementary File for details) to solve optimization problems with large number of parameters.

In the context of IBCGA, each individual will be called a chromosome containing genes. Each of the biophysico properties works as a binary gene constructing the chromosome. A value of 1 (0) for any binary gene means it is being considered (not considered) for the prediction of SLR. The feature space is constructed by taking the biophysico properties having value 1 and concatenating the values of these properties for all of the amino acids in the *context window*. The centered lysine residue is excluded in this step because it is the same across all the samples. An RF is trained and the achieved MCC of the classifier on the corresponding validation dataset works as the fitness of that individual. The detailed steps to calculate the fitness of an individual is demonstrated in Figure 1.

At each iteration, the IBCGA algorithm maintains a number of solutions each having *r* genes with value 1, where *r*_*start*_ ≤ *r* ≤ *r*_*end*_. Here, *r* is the number of 1’s in each chromosome. The steps of the algorithm with the given values of *r*_*start*_ and *r*_*end*_ are as follows [9].

Step 1. Generate *N*_*pop*_ random individuals (chromosomes) each having *r* genes with value 1.

Step 2. Evaluate the fitness of the individuals using the procedure described in Figure 1.

Step 3. Select *P*_*c*_ × *N*_*pop*_ individuals from the current population by the 2-way tournament selection algorithm in pairs and perform orthogonal array crossover on each of these pairs. Here, *P*_*c*_ is the crossover probability.

Step 4. Two genes’ values are swapped as part of mutation for *P*_*m*_ × *N*_*pop*_ individuals. This mutation is not applied on the best individual for any specific *r* in order to preserve the best fitness value.

Step 5. Repeat Steps 2 to 4 *N*_*iter*_ times. In the (*N*_*iter*_ + 1)*th* time, go to Step 6.

Step 6. Randomly change one binary gene’s value from 0 to 1 for each of the *N*_*pop*_ individuals in order to increase the value of *r* by 1. If *r* ≤ *r*_*end*_ go to Step 2. Else, terminate the algorithm.

### 2.5 Deep Learning Architecures

#### 2.5.1 Model Overview

We consider only simpler architectures with lesser parameters (Table 2). We initially experiment with two different architectures, where we leverage the power of CNN and Bi-LSTM architectures. Our two basic settings are **Bi-LSTM+CNN** (*BLC*) and **CNN+Bi-LSTM** (*CBL*). As the name indicates, the main difference between BLC and CBL models lies in the order the two constituent deep neural network architectures have been connected to each other. We then use the combination of these two architectures to build a better predictor (referred to as *CBL_BLC* henceforth).

**Table 2.** Number of trainable parameters in each of the deep learning architectures.

| Model | CBL | BLC | CBL_BLC |
| --- | --- | --- | --- |
| Trainable parameters | 5489 | 8465 | 19025 |

The reason for choosing CNN and Bi-LSTM as our fundamental models is not arbitrary. As protein sequence is a sequential data, both 1D-CNN and LSTM can extract features from the protein sequence. However, CNN’s main function is to extract local features whereas LSTM can capture the long-range dependency in the sequence. Notably, Zhang *et al*. experimented with CNN-LSTM, LSTM-CNN architectures in [29] for building SLR predictors. The architectures of *CBL, BLC* and *CBL_BLC* are shown in Figure 2. The *None* in each cell represents the *batch size*.

**Fig. 2.**
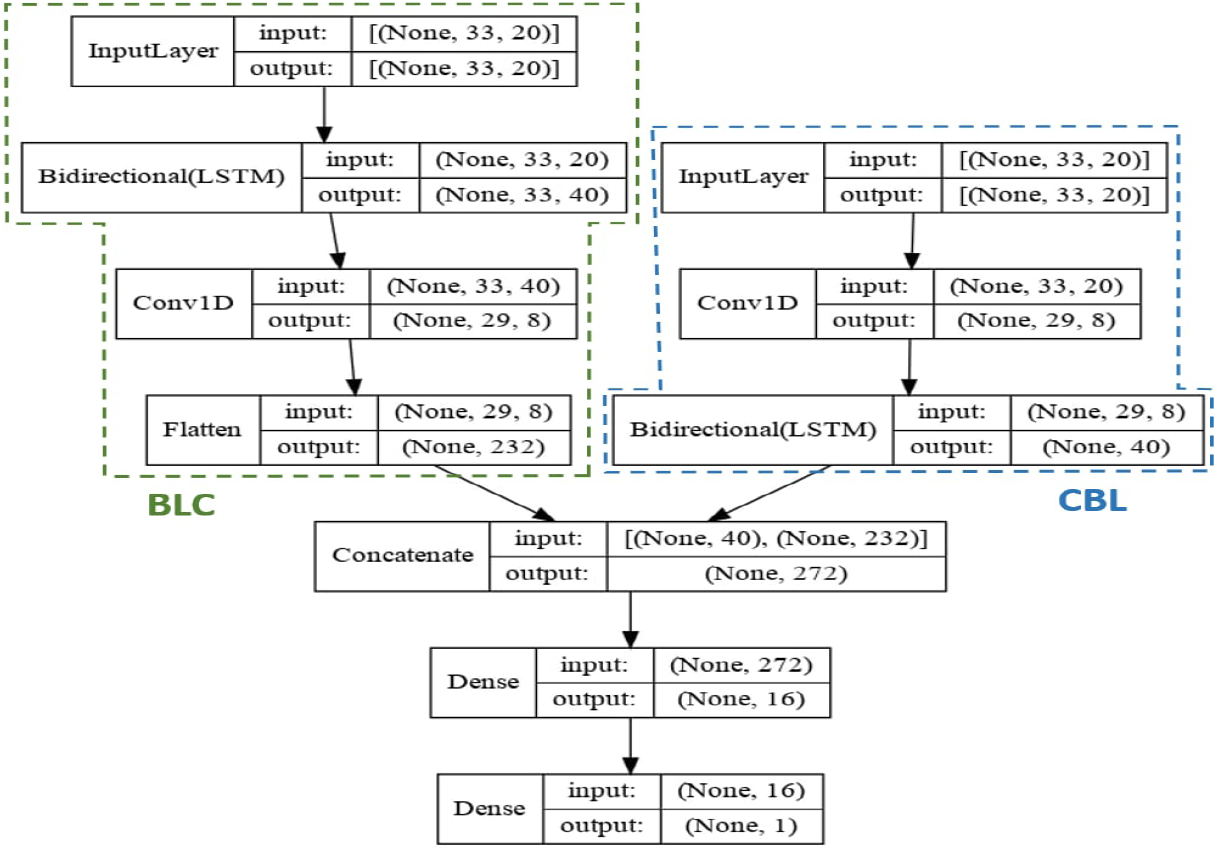
The CBL_BLC architecture has two branches. The left branch corresponds to the architecture of BLC. The one-hot encoded (details of one-hot encoding is discussed in Section 4 of Supplementary File) protein sequences are first fed into bidirectional LSTM layer. 1D-CNN layer follows this layer. The output of the CNN layer goes through a flatten layer. The right branch corresponds to the architecture of CBL. Here, the one-hot encoded protein sequences are first fed into 1D convolution layer. The local features extracted from the CNN layer are then fed into the Bidirectional LSTM layer. In CBL_BLC, the features from these two architectures are concatenated and passed through two densely connected layers.

#### 2.5.2 Loss functions and Checkpoints

We experimented with three loss functions (appropriate for binary classification), namely, binary cross-entropy, hinge loss and squared hinge loss (see Section 5 of Supplementary File for details of these loss functions) and choose binary cross-entropy as this gives better results than the other two. During training the deep learning architectures with **D2** dataset, we use *weighted binary cross-entropy* (see Section 5.1 of Supplementary File) because the number of negative samples is twice the number of positive samples. So, we penalize the model two times more for predicting a 1 as 0 than predicting a 0 as 1 by setting (*w*_0_, *w*_1_) = (1, 2). We run our model with *batch size* set to 128 and for 80 epochs. We monitor the loss on the validation dataset after each epoch and save a checkpoint of the model if the calculated loss is smaller than the smallest loss found so far. After the training is complete, we use the model that has had the smallest validation loss across the 80 epochs.

#### 2.5.3 Ensembling of different models

We train each of the three architectures for 5 times. This results in 5 different versions for each architecture. We calculate the average probability of a sample’s belonging to class 1 according to the following equation:

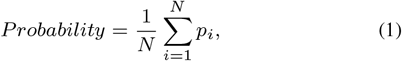

where, *p*_*i*_ is the predicted probability from the *i*^*th*^ classifier. If this probability is less than a pre-defined *threshold*, the ensemble classifier classifies the sample as 0, otherwise the sample is classified as 1. We initially set the *threshold* value to 0.5 to evaluate the performance of the ensemble classifiers.

#### 2.5.4 Tuning the threshold for ensemble classifiers

As has been discussed in Section 2.5.3, we have a parameter called *threshold* which is initially set to 0.5. However, if we decrease (increase) the threshold value, the SN (SP) will increase compromising the SP (SN). Hence, this parameter can be tuned to produce ‘better’ results under different circumstances where it may be more desirable to achieve a better SP or SN for that matter. We use differential evolution algorithm (see Section 6 of Supplementary File for details) to find an optimal value for the threshold to improve the performance. Each individual in this algorithm refers to a threshold value. The fitness of an individual is the MCC value obtained by the ensemble classifier on the validation dataset.

## 3 Results and Discussions

### 3.1 Performance of combination of biophysico properties

The 29 properties derived from Section 2.3 is the universal set of properties for the IBCGA algorithm. We set *r*_*start*_ = 1, *r*_*end*_ = 20, *N*_*pop*_ = 50, *N*_*iter*_ = 5. We take the values of both *P*_*m*_ and *P*_*c*_ from the set {0.5, 0.6, 0.7, 0.8, 0.9} resulting in 25 different combinations of *P*_*m*_ and *P*_*c*_. The best MCC values obtained on both the validation datasets of D1 and D2 for each value of *r* is shown in Figure 3.

**Fig. 3.**
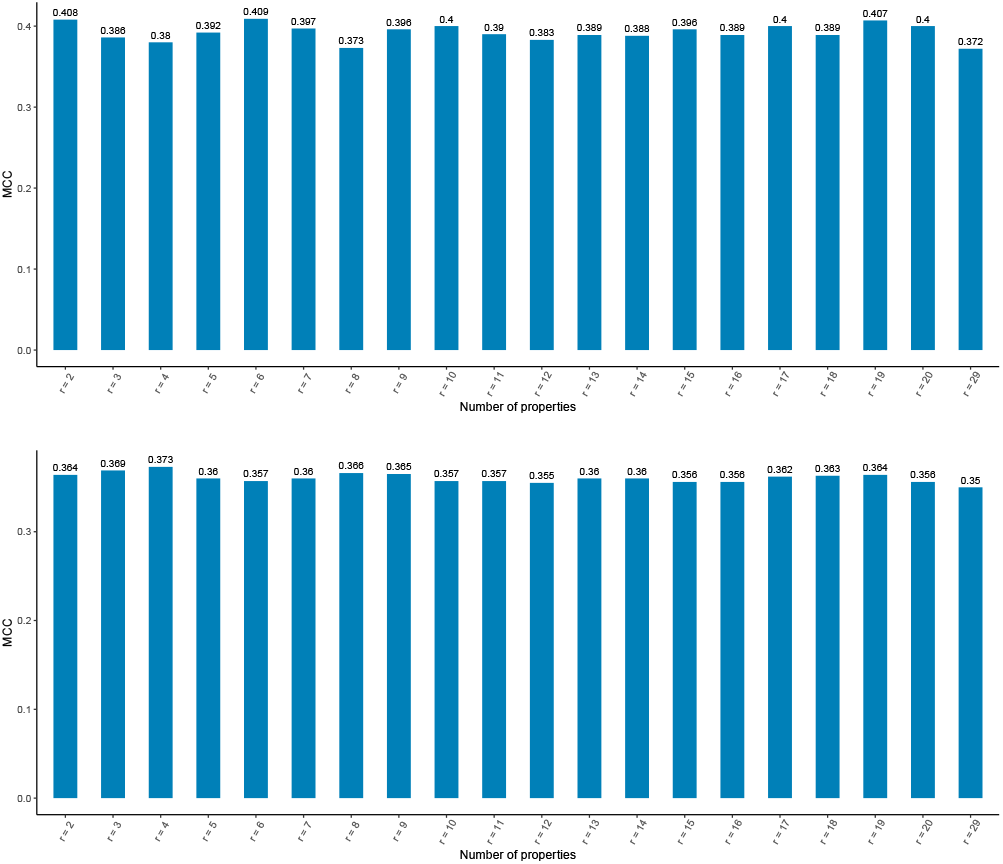
The IBCGA algorithm was run for 25 different pairs of values for *P*_*c*_ and *P*_*m*_. The best MCC value obtained among the 25 values for each value of *r* is recorded. These are the histogram plots of the best MCC values for D1 and D2 against each value of *r* from 2 to 20. *r* = 29 means we are using all the 29 biophysico properties.

We observe that using all the top 29 biophysico properties is giving us comparatively poorer performance if compared with the performances of *r* = 2 to *r* = 20. This proves the necessity of searching for suitable combinations rather than using all the better performing properties as has been done by several works in the literature [7, 6, 16, 29]. Hence, we obtain 19 models trained by RF for *r* = 2 to *r* = 20 for both D1 and D2 datasets.

### 3.2 Performance of deep learning architectures

We use Adam optimizer to train CBL, BLC and CBL_BLC. We perform the training of each of the architectures 5 times and report the average performance on the validation dataset in Figure 4. We observe that the CBL_BLC is the winner with respect to SP, ACC & MCC for both datasets. CBL achieves the highest SN for both datasets at the cost of the lowest SP. From SN-SP trade-off point of view, CBL_BLC is the best for D1 and BLC is the best for D2 because they achieve very good SN without compromising SP.

**Fig. 4.**
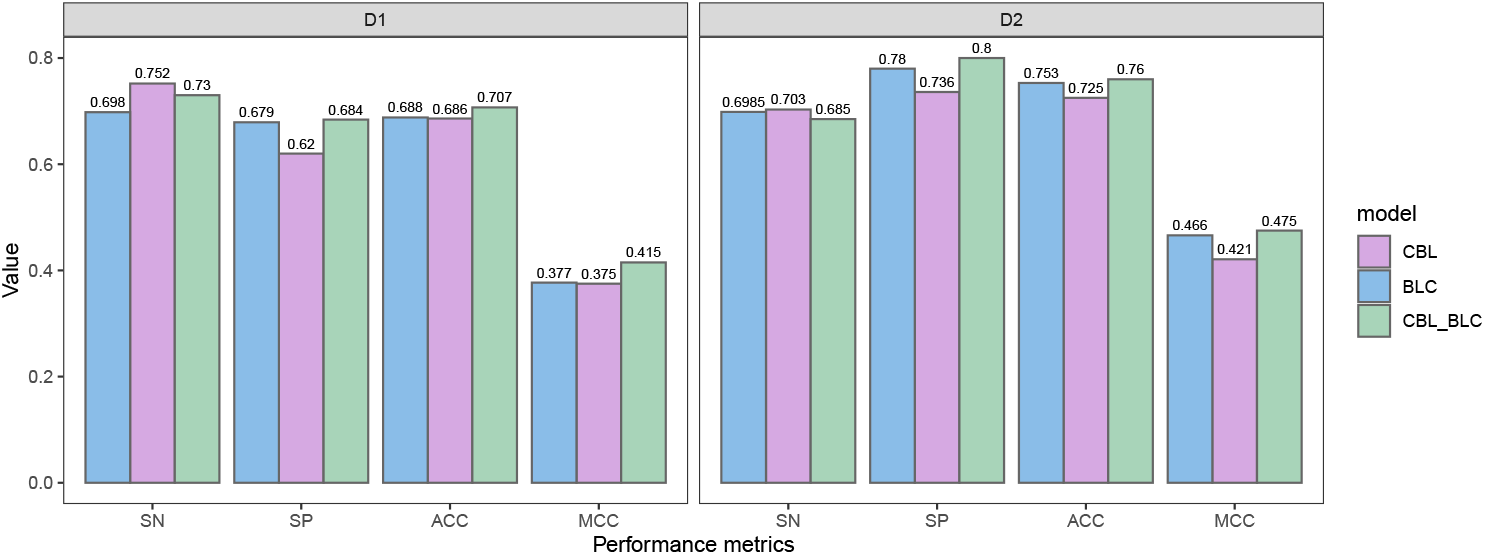
Performance of CBL, BLC and CBL_BLC on the validation dataset of D1 and D2. Five independent runs are conducted for each architecture and dataset combination, and the average performance values are reported. The SN of CBL is the best for both datasets. For the other three metrics, CBL_BLC is the winner for both datasets.

### 3.3 Performance of ensemble classifiers

As has been already mentioned in Section 2.5.3, we have run each of the models for 5 times. We can ensemble these 5 models as we expect that even if some of the 5 models misclassify a sample, majority will correctly classify that one. We have obtained 19 models from the combination of biophysico properties (see Section 3.1). We will refer to the collection of these models as *BP*. We denote the ensemble classifier of any architecture by appending ‘-E’ to the respective name. Hence, CBL-E will represent the ensemble classifier of CBL and so on. Similarly, the ensemble of CBL and BLC classifiers will be denoted as (CBL+BLC)-E. The performance of the ensemble classifiers are shown in Table 3.

**Table 3.** Performance of different ensemble classifiers on the validation dataset of D1 and D2. (CBL+BLC+CBL_BLC+BP)-E achieves best SN, ACC, MCC in D1. (CBL+BLC)-E achieves best SN in D2. (BLC+CBL_BLC)-E achieves best SP in D1. CBL_BLC-E achieves best SP, ACC, MCC in D2.

| Architecture | D1 |  |  |  | D2 |  |  |  |
| --- | --- | --- | --- | --- | --- | --- | --- | --- |
|  | SN | SP | ACC | MCC | SN | SP | ACC | MCC |
| CBL-E | 0.764 | 0.624 | 0.695 | 0.392 | 0.702 | 0.744 | 0.73 | 0.429 |
| BLC-E | 0.717 | 0.685 | 0.701 | 0.402 | 0.712 | 0.791 | 0.765 | 0.49 |
| CBL_BLC-E | 0.744 | 0.691 | 0.718 | 0.436 | 0.6985 | **0.81** | **0.772** | **0.498** |
| BP-E | 0.745 | 0.64 | 0.695 | 0.391 | 0.721 | 0.66 | 0.681 | 0.361 |
| (CBL+BLC)-E | 0.759 | 0.657 | 0.708 | 0.418 | **0.724** | 0.775 | 0.758 | 0.483 |
| (BLC+CBL_BLC)-E | 0.7299 | **0.692** | 0.711 | 0.423 | 0.707 | 0.8 | 0.769 | 0.496 |
| (CBL+BLC+CBL_BLC)-E | 0.748 | 0.678 | 0.713 | 0.428 | 0.717 | 0.787 | 0.764 | 0.49 |
| (CBL+BLC+CBL_BLC+BP)-E | **0.772** | 0.681 | **0.726** | **0.454** | 0.723 | 0.766 | 0.752 | 0.472 |

CBL_BLC-E dominates CBL-E and BLC-E on both D1 and D2 datasets except with respect to SN. In case of (CBL+BLC)-E, (BLC+CBL_BLC)-E and (CBL+BLC+CBL_BLC)-E, the performances are not improving much from the individual ensemble classifiers (i.e., CBL-E, BLC-E and CBL_BLC-E). If we observe the performance of BP-E both on D1 and D2 datasets, we note that the SN (0.745 on validation set of D1, 0.705 on validation set of D2) is competitive if compared to the SNs of CBL-E, BLC-E and CBL_BLC-E. However, BP-E is lagging way behind with respect to SP which makes the other metrics (ACC and MCC) poor too. However, we achieve the highest SN, ACC and MCC on the dataset D1 with (CBL+BLC+CBL_BLC+BP)-E which shows the usefulness of the suitable combinations of biophysico properties. Although (CBL+BLC+CBL_BLC+BP)-E is not the best with respect to any metric on the dataset D2, the SN is almost touching the highest value (i.e., 0.724).

### 3.4 Tuning the threshold parameter

We use the differential evolution class from the scipy.optimize (scipy version 1.2.0) package to optimize the MCC value. We keep the lower bound of the *threshold* as 0.4 and upper bound as 0.6. The *tol* (tolerance) parameter is set to 1e-7. Other parameters are kept as the default ones. The algorithm is run 10 times and the threshold value that gives the best MCC value is recorded. The comparison of the tuned models along with the untuned models are shown in Figure 5. We observe that with the tuned threshold values, the ensemble classifiers are able to achieve better MCC compared to the MCC achieved by the ensemble classifiers with default threshold (i.e., 0.5).

**Fig. 5.**
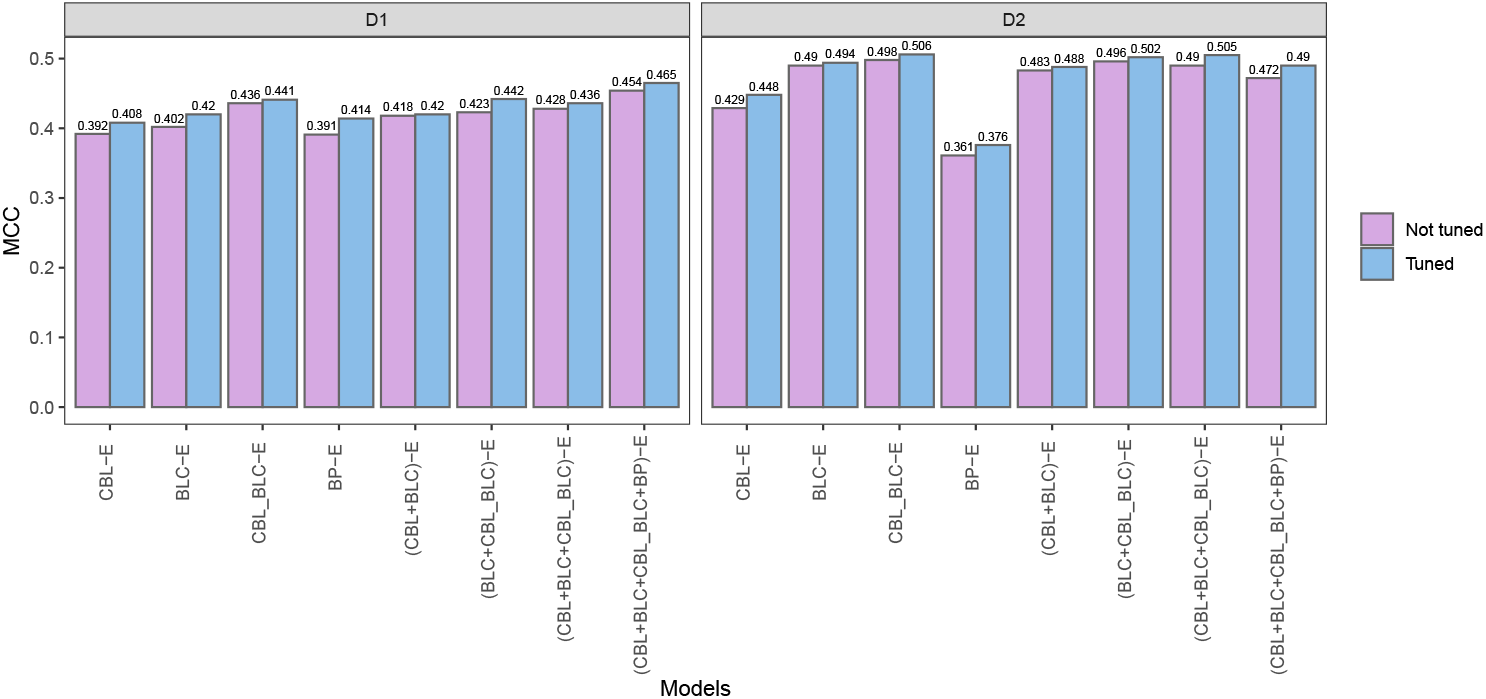
Performance of tuned and untuned ensemble classifiers. For all the models on both datasets, we see an improvement in MCC with the tuned value compared to the untuned value of the threshold.

### 3.5 Comparison with existing works

We choose the (CBL+BLC+CBL_BLC+BP)-E for dataset D1 to compare with the existing works as this achieves the best MCC value. For dataset D2 we consider both CBL_BLC-E and (CBL+BLC+CBL_BLC)-E models for the following two reasons: (a) both are quite competitive with each other and (b) although second best, (CBL+BLC+CBL_BLC)-E is the ensemble of higher number of classifiers. Hence, the probability of the unseen samples from the test dataset to be correctly classified is higher with (CBL+BLC+CBL_BLC)-E.

Note that, if the comparison is being conducted on the same dataset (i.e., both training and test datasets are same), we directly use the reported results from the respective paper. In the case when we have to compute the results for a predictor on a different dataset, we re-implement the predictor and train with the same training data that is being used to train our model. The training and testing dataset of D1 has been directly used by LSTMCNNsucc [10]. Although DeepSuccinylSite [22] used solely the training and testing dataset of D2, we re-implemented this predictor to re-calculate the performance of DeepSuccinylSite because it undersampled the test dataset before evaluation but other previous works didn’t. Hence, we are able to calculate its performance on D1 too. The results of these predictors along with the performance of both (CBL+BLC+CBL_BLC+BP)-E and tuned (CBL+BLC+CBL_BLC+BP)-E on the test dataset of D1 are shown in Table 4. We observe that both of our models are performing better than DeepSuccinylSite. However, LSTMCNNSucc has a significantly better SP at the cost of a very poor SN indicating that this method is performing poorly on the actual task, which is to predict the positive sites as positive.

**Table 4.** Performance of some existing works on the testing dataset of D1 along with the performance of (CBL+BLC+CBL_BLC+BP)-E (with threshold=0.5 and tuned threshold)

| Method | Threshold | SN | SP | ACC | MCC | Remarks |
| --- | --- | --- | --- | --- | --- | --- |
| LSTMCNNSucc | - | 0.592 | **0.796** | **0.779** | **0.251** | Low SN |
| DeepSuccinylSite | - | 0.725 | 0.593 | 0.604 | 0.176 | Low SP |
| (CBL+BLC+CBL_BLC+BP)-E | 0.5 | 0.751 | 0.677 | 0.683 | 0.246 | High SN |
| Tuned (CBL+BLC+CBL_BLC+BP)-E | 0.491 | **0.769** | 0.658 | 0.668 | 0.24 | Highest SN, low SP |

The tools Succinsite, SuccinSite2.0, GpSuc, Psucce, Inspector, DeepSuccinylSite and DeepSucc used the training and testing dataset of D2. The results of these predictors along with the performance of (CBL+BLC+CBL_BLC)-E (with different thresholds) on the test dataset of D2 are shown in Table 5. We observe that (CBL+BLC+CBL_BLC)-E with a threshold of 0.5 achieves a moderate SN and SP and as a result, has a higher MCC value than the MCC value of all other methods except the MCC achieved by DeepSucc. Unfortunately, as is marked in Table 5, the results reported in [29] is unreliable at best; Section 3.6 discusses several issues with this predictor.

**Table 5.**
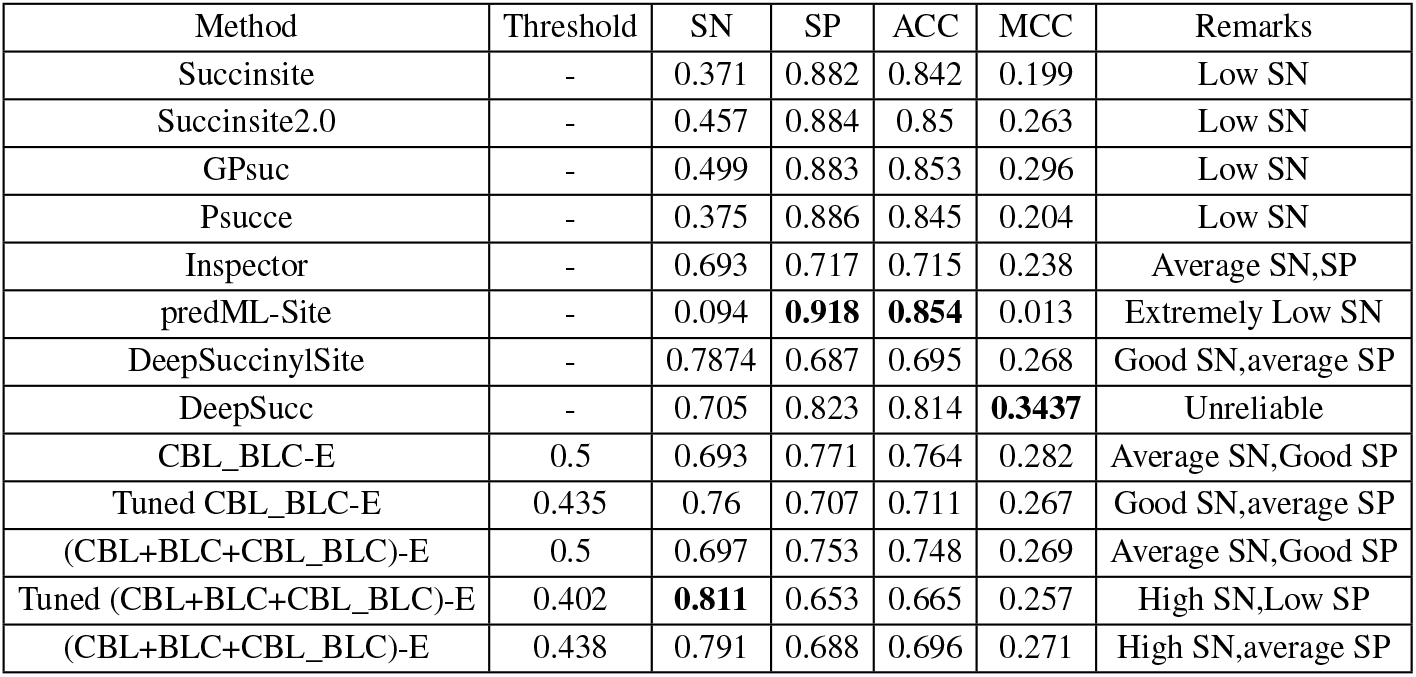
Performance of several existing works on the testing dataset of D2 along with the performance of CBL_BLC-E (with threshold=0.5, tuned threshold) and (CBL+BLC+CBL_BLC)-E (with threshold=0.5, tuned threshold and for another example threshold, namely, 0.438)

Tuned (CBL+BLC+CBL_BLC)-E achieves the highest SN among the existing works at the cost of a low SP. CBL_BLC-E achieves a better SP and MCC compared to the SP and MCC of (CBL+BLC+CBL_BLC)-E. However, the tuned CBL_BLC-E does not achieve a good SN if compared with the SN of tuned (CBL+BLC+CBL_BLC)-E. It is evident from the result of tuned (CBL+BLC+CBL_BLC)-E that if we increase the threshold a little from the tuned value, SN will decrease and SP will increase. We have shown an example in Table 5 with a threshold value that produces better result than DeepSuccinylSite with respect to both SN and SP. Although unrealiable, we have to note that the performance of DeepSucc is better in all cases due to a very high SP at the cost of a moderate SN. We will present a more detailed discussion on this method in the following section.

### 3.6 A Discussion on DeepSucc [29]

During the write-up stage of this paper, the publication of a new predictor, called DeepSucc [29] came to our knowledge. We thoroughly investigated the codebase shared by the paper and found some issues which are discussed below.

- They performed cross-validation to compare among different architectures. But in their code, we find that after each fold of cross-validation was performed, the authors mistakenly did not re-initialize the variables which has resulted in a data leakage. Hence, the claimed results are completely unreliable. And our repeated communications with the authors did not get any response.
- The code which was used for performing testing on the dataset D2 is completely absent in the repository. Although there is a folder named “Test” inside the repository, all the codes correspond to the cross-validation codes.
- We re-implemented their architectures from the provided code removing the errors but the produced results were far worse than the claimed ones.

For the above-mentioned reasons, the DeepSucc’s results seem unreliable at best. Therefore, although we have reported the results of DeepSucc (as reported in their paper), we actually exclude those from our comparative analysis and discussion.

### 3.7 SN-SP tradeoff

It is evident from earlier discussion that increasing the threshold will increase (decrease) the SP (SN) and vice versa. The SN and SP of three of our ensemble classifiers for different threshold values ranging from 0.4 to 0.6 along with the SN and SP of some of the notable existing classifiers are shown in Figure 6.

**Fig. 6.**
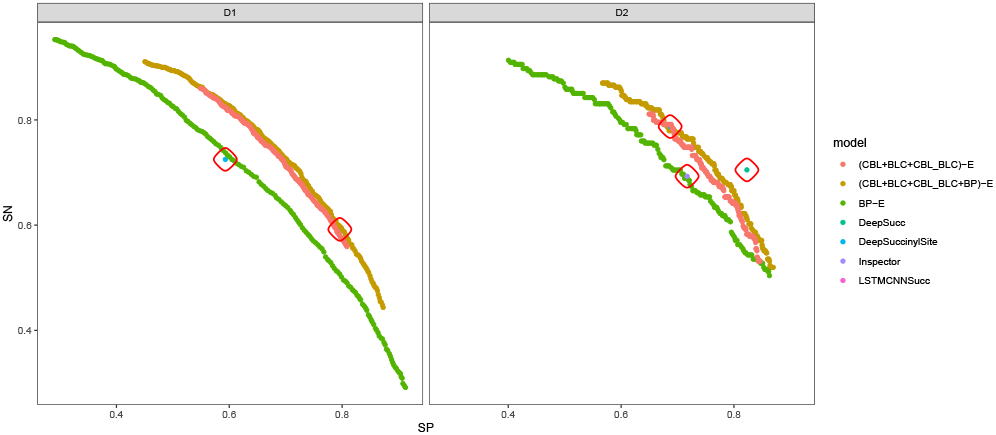
SN vs SP of several ensemble classifiers and of some notable existing predictors for different threshold values ranging from 0.4 to 0.6 for both D1 and D2. The circles focus the existing classifiers.

We observe that our best classifiers (i.e., (CBL+BLC+CBL_BLC+BP)-E for D1, (CBL+BLC+CBL_BLC)-E for D2) are better than all of the existing predictors. Although BP-E is lagging far behind the other two classifiers for both datasets, we can see a sharper increase in SP and sharper decrease in SN as the threshold increases for BP-E compared to the other two classifiers.

## 4 Conclusion

In this paper, we have proposed an optimization algorithm to search for suitable combinations of biophysico properties for better prediction of succinylated lysine residues. We have achieved better performance with a few properties compared to the performance achieved through combining all the top performing properties. We have experimented with some simple deep learning architectures CBL, BLC and CBL_BLC with lesser number of trainable parameters. We have also employed different ensembling techniques to improve upon the performance of our models, which included heterogeneous ensembling of traditional ML models with deep learning architectures as well. Finally, we have applied differential evolution to tune the threshold of ensemble classifiers thereby providing the biologists and practitioners with a knob to balance the SN and SP. We have showed that our models have achieved better results than the existing state-of-the-arts through varying the threshold value. We believe that our methodology, if applied on state-of-the-art giant models like XLNet [28], MPNet [19] will produce even better results, albeit at the cost of huge money, energy and time which is against the concept of GreenAI [18]. Also, the models will have to be re-trained for different datasets which would be infeasible in most cases. The main motivation of constructing computational models for detecting PTM sites is that the wet lab experimental methods are costly and time-consuming. If the proposed computational model is also time consuming and costly to train, the purpose is not served. We believe that our simple deep learning models along with the suitable combinations of biophysico properties will serve as a useful tool for the biologists to filter the lysine residues on which further wet lab experiments will be conducted.

## Supplementary information

Supplementary data are available at https://github.com/Dariwala/Succinylation-with-biophysico-and-deep-learning/Supplementary Information

## Notes

https://github.com/Dariwala/Succinylation-with-biophysico-and-deep-learning

### Competing Interest Statement

The authors have declared no competing interest.

