## Supplementary File for "Succinylated lysine residue prediction revisited"

#### 1 Preprocessing Datasets

### 1.1 D1

Succinylated proteins (6377) are collected from the Protein Lysine Modification Database (PLMD) [8]. To remove proteins having above threshold sequence similarity, we apply CD-Hit with cutoff set to 40%. Removing homology, we obtain 3560 proteins. The experimentally verified lysine residues are considered as the positive sites, and all other lysine residues in the same proteins are considered as the negative sites. We randomly divide the proteins into training and test set in the ratio 4:1. There are 6812 succinylated lysine residues in the training set. But, the number of non-succinylated lysine residues is more than 9 times higher. We randomly discard negative samples in order to make the training dataset balanced. Thus, in the processed training dataset, we have 6812 non-succinylated lysine residues from 2848 proteins. We separate out 300 proteins from the training set as the validation set for tuning several hyper-parameters. So, the final training set contains 2548 proteins.

### 1.2 D2

We use another datatset for comparative analysis which is compiled by [1] as follows. At first, 10,000 succinylated proteins are collected from nine species. Then, redundancy is removed by applying CD-HIT with cutoff set to 30%. The training dataset contains 2,198 proteins with 4,750 succinylated and 9,500 non-succinylated lysine residues. We randomly separate out 3000 samples from the training set for tuning several hyper-parameters, of which 960 are succinylated and 2040 are non-succinylated lysine residues. The test dataset has 124 proteins with 254 succinylated and 2,977 non-succinylated lysine residues.

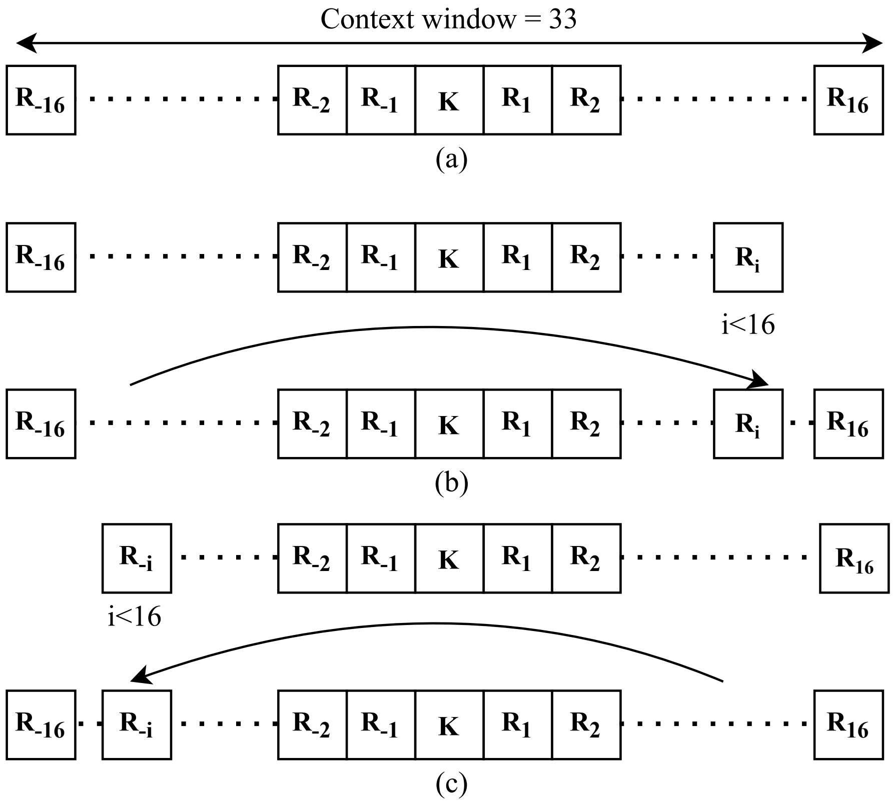


Figure 1: Creating positive and negative samples with the neighboring amino acids of the lysine residue. (a) The lysine residue has enough neighbors both on its downstream and upstream. (b) The lysine residue lacks neighbors on its downstream. So, mirror effect is used to bring amino acids from its upstream. (c) The lysine residue lacks neighbors on its upstream. So, mirror effect is used to bring amino acids from its downstream.

#### 2 Performance Metrics

Some popular metrics for binary classification problems are Sensitivity (SN), Specificity (SP), Accuracy (ACC), Matthews Correlation Coefficient (MCC).

The equations to determine these metrics are as follows.

$$Sensitivity\ (SN) = \frac{TP}{TP + FN}$$

$$Specificity\ (SP) = \frac{TN}{TN + FP}$$

$$Accuracy\ (ACC) = \frac{TP + TN}{TP + TN + FP + FP}$$

$$MCC = \frac{(TP \times TN) - (FP \times FN)}{\sqrt{(TP + FN) \times (TN + FP) \times (TP + FP) \times (TN + FN)}}$$

Here, $TP$, $TN$, $FP$, and $FN$ refers to True Positive, True Negative, False Positive, and False Negative, respectively.

### 3 Orthogonal Experiment Design

Orthogonal experimental design (OED) with orthogonal array (OA) is an effective way of analyzing the effect of several factors simultaneously [7]. The factors work as parameters, which affect the response variables. The factors can take different values known as levels of the factors. A **complete factorial** experiment involves trying all possible combinations of levels of each of the factors which is often infeasible [6]. A **fractional factorial** experiment tries to select a subset of combinations of levels of the factors. Orthogonal experimental design uses fractional factorial experiments to efficiently determine the best combination of factor levels to optimize the response variable. An illustrative example of OED using an objective function is as follows:

$$maximize\ y(x_1, x_2, x_3) = 5x_1 - 2x_2 + x_3$$

where $x_1 \epsilon \{-1, 1\}$, $x_2 \epsilon \{1, 2\}$ and $x_3 \epsilon \{3, 4\}$.

This maximization problem can be considered as an experimental design problem of three factors, with two levels each. Let, $x_1$, $x_2$, $x_3$ be factors $F_1$, $F_2$, and $F_3$, respectively. Let the smaller (larger) value of each parameter be the level 1 (level 2) for each factor. The objective function is the response variable, $y$. A **complete factorial** experiment would evaluate $2^3 = 8$ level combinations. As a result, the best combination $(x_1, x_2, x_3) = (1, 1, 4)$ with $y = 7$ is obtained. The results of the complete factorial experiment are shown in Table 1.

A fractional factorial experiment uses a well-balanced subset of level combi-

Table 1: An example of complete factorial experiment

| Combination No. | Factors | | | Parameters | | | Response variable |
|---|---|---|---|---|---|---|---|
| | $F_1$ | $F_2$ | $F_3$ | $x_1$ | $x_2$ | $x_3$ | |
| 1 | 1 | 1 | 1 | -1 | 1 | 3 | -4 |
| 2 | 1 | 1 | 2 | -1 | 1 | 4 | -3 |
| 3 | 1 | 2 | 1 | -1 | 2 | 3 | -6 |
| 4 | 1 | 2 | 2 | -1 | 2 | 4 | -5 |
| 5 | 2 | 1 | 1 | 1 | 1 | 3 | 6 |
| 6 | 2 | 1 | 2 | 1 | 1 | 4 | 7 |
| 7 | 2 | 2 | 1 | 1 | 2 | 3 | 4 |
| 8 | 2 | 2 | 2 | 1 | 2 | 4 | 5 |

nations, such as the second, fourth, fifth, and seventh combinations. The best one of the four combinations is $(x_1, x_2, x_3) = (1, 1, 3)$ with $y = 6$. Using OA, the best combination $(x_1, x_2, x_3) = (1, 1, 4)$ can be obtained by judiciously selecting a subset of the combinations. The procedure of constructing OA is described below.

Let, $N$ be the number of factors, each of which has two levels. Thus, a complete factorial experiment will include trying out $2^N$ combinations. To construct an OA with $N$ factors, let, $n = 2^{\lceil log_2(N+1) \rceil}$. Now, build a two level OA, $L_n(2^{n-1})$, with $n$ rows and $n-1$ columns. Only the first $N$ columns of this table will be used. The numbers 1 and 2 in each column indicate the levels of the factors. Each column has an equal number of 1s and 2s. Each of the four pairs $(1, 1)$, $(1, 2)$, $(2, 1)$ and $(2, 2)$ will appear same number of times in each possible pair of the $n-1$ columns. For example, an $L_4(2^3)$ is as follows.

| | | |
|---|---|---|
| 1 | 1 | 1 |
| 1 | 2 | 2 |
| 2 | 1 | 2 |
| 2 | 2 | 1 |

The algorithm used to generate OAs is described in Algorithm 1 (taken from [2]).

#### 3.1 Factor analysis with Orthogonal Array

Factor analysis is conducted to compute the effect of each factor on the objective function, rank the most effective factors, and determine the best best

**Algorithm 1** Creating Orthogonal Array OA

**Require:** $N \geq 1$

$n \leftarrow 2^{\lceil \log_2 N+1 \rceil}$<br>
**for** $i$ *from* 1 *to* $N$ **do**<br>
    **for** $j$ *from* 1 *to* $N$ **do**<br>
        $level \leftarrow 0$<br>
        $k \leftarrow 0$<br>
        $mask \leftarrow \frac{n}{2}$<br>
        **while** $k \geq 0$ **do**<br>
            **if** $k\%2 \neq 0$ & $bitwise\_AND(i-1, mask) \neq 0$ **then**<br>
                $level \leftarrow (level+1)\%2$<br>
            **end if**<br>
            $k \leftarrow \lfloor \frac{k}{2} \rfloor$<br>
            $mask \leftarrow \frac{mask}{2}$<br>
        **end while**<br>
        $OA[i][j] \leftarrow level + 1$<br>
    **end for**<br>
**end for**

level for each factor in order to optimize the objective function. An illustrative example of orthogonal experiment design using a 2-level orthogonal array $L_M(2^{M-1})$ with $M$ rows and $M-1$ columns is shown in Table 2. In this example, $M=4$, there are three factors each of which having two levels (1 or 2). 1 corresponds to the exclusion and 2 corresponds to the inclusion of the factor in the proposed feature selection. Let $f_t$ denote the value of the objective function for combination $t$. Define the main effect of factor $j$ with level $k$ as $S_{jk}$ where $j = 1, 2, 3, \ldots, M-1$ and $k = 1, 2$ as:

$$S_{jk} = \sum f_t \cdot F_t \tag{1}$$

where $F_t = 1$ if the level of factor $j$ of combination $t$ is $k$; otherwise, $F_t = 0$. Since the objective function is to be maximized, the level 1 of factor $j$ makes a better contribution to the function than level 2 of factor $j$ does when $S_{j1} > S_{j2}$. After the better of the two levels for each factor is determined, a good combination consisting of all factors with the better levels can be easily reasoned.

The Rank column in Table 2 shows the rank of the combination $t$ among all possible ($2^3 = 8$) combinations. In this example, the reasoned combination gets the best value for the objetive function which may not be the case in practical life.

Table 2: A sample orthogonal experiment

| $t$ | **Factors** | | | $f_t$ | **Rank** |
|---|---|---|---|---|---|
| | 1 | 2 | 3 | | |
| 1 | 1 | 1 | 1 | 35.3% | 6/8 |
| 2 | 1 | 2 | 2 | 32.1% | 8/8 |
| 3 | 2 | 1 | 2 | 60.5% | 2/8 |
| 4 | 2 | 2 | 1 | 57.8% | 3/8 |
| $S_{j1}$ | 67.4 | 95.8 | 93.1 | | |
| $S_{j2}$ | 118.3 | 89.9 | 92.6 | | |
| MED | 50.9 | 5.9 | 0.5 | | |
| Rank | 1 | 2 | 3 | | |
| Better level | 2 | 1 | 1 | 63.2% | 1/8 |

#### 3.2 Orthogonal Array Crossover

In our context, each sample $S_i$ is a vector consisting of binary genes each of which represents a biophysico property. 1 (0) means we will consider (not consider) that property for assessing the fitness of the individual. Let's introduce some notations for ease of explanation of the algorithm.

- If $S = \{s_i \mid 1 \leq i \leq n\}$, then $S[p : q] = \{s_i \mid p \leq i \leq q\}$.
- $\zeta(S)$ denote the number of 1's in $S$.
- The cut point $C = \{c_i \mid 1 \leq i \leq n\}$ of two individuals $S_1, S_2$ is defined such that for each element $c_i$ of $C$,
  - $S_1[c_{i-1} : c_i] \neq S_2[c_{i-1} : c_i]$
  - $\zeta(S_1[c_{i-1} : c_i]) = \zeta(S_2[c_{i-1} : c_i])$ (if $i = 1$, then $c_{i-1} = 1$)

  And, no other set with cardinality greater than the cardinality of $C$ has the same properties.

During crossover of two samples $S_1$ and $S_2$, we first calculate their cut point $C$. We consider each element $c_i$ of $C$ as a factor with two levels 1 and 2. 1 means we will consider $S_1[c_{i-1} : c_i]$ to create the child, 2 means we will consider $S_2[c_{i-1} : c_i]$ to create the child. We then create orthogonal array and perform orthogonal experiment and find out the best level combination. The best level combination gives us one child. The second child is created by toggling the value of the worst factor.

#### 4 One-hot Encoding

The samples are first pre-processed according to the procedure mentioned in Supplementary "Section 1". Then, each amino acid is mapped to a unique value ranging from 0 to 19 (as there are 20 possible amino acids). After that, each integer value is converted into a 20 dimensional vector with all zeros except a 1 at that integer valued index. For example, if the lysine residue 'K' is mapped to the integer 11, the final 20 dimensional vector from the lysine residue will be $\{0, 0, 0, 0, 0, 0, 0, 0, 0, 0, 1, 0, 0, 0, 0, 0, 0, 0, 0, 0\}$. Hence, if we use *context window* = 33, the dimension of input to the deep learning model will be $33 \times 20$.

#### 5 Loss function

In the context of an optimization algorithm, the function used to evaluate a candidate solution (i.e., a set of weights) is referred to as the objective function. Typically, with neural networks, we want to minimize the error. As such, the objective function is referred to as the loss function.

The three most popular loss functions for binary classifications are:

**Binary Cross-Entropy Loss** Let, there are $N$ samples in a dataset and $y_i$ denote the label (0 or 1) of the $i^{th}$ sample. Then, binary cross-entropy loss of these $N$ samples is calculated as follows:

$$Loss = -\frac{1}{N}\sum_{i=1}^{N}(y_i \log p_i + (1 - y_i)\log(1 - p_i)), \tag{2}$$

where, $p_i$ is the predicted probability of the $i^{th}$ sample to belong to class 1.

**Hinge Loss** For computation of hinge loss, the outputs must be in the set $\{-1, 1\}$. For an intended output $t = \pm 1$ and a classifier score $y$, the hinge loss of the prediction $y$ is defined as:

$$l(y) = max(0, 1 - t \cdot y) \tag{3}$$

**Squared Hinge Loss** Squared hinge loss has the effect of the smoothing the surface of the error function and making it numerically easier to work with. When the hinge loss requires better performance on a given binary classification problem it is mostly observed that a squared hinge loss may be appropriate to use. For computation of squared hinge loss, the outputs must be in the set $\{-1, 1\}$. For an intended output $t = \pm 1$ and a classifier score $y$, the squared hinge loss of the prediction $y$ is defined as:

$$l(y) = max(0, 1 - t \cdot y)^2 \tag{4}$$

##### 5.1 Weighted Binary Cross-entropy

The **weighted binary cross-entropy** is used when we want to give more penalty to the model for misclassifying samples of a specific class. The formula for weighted binary cross-entropy is as follows:

$$Loss = -\frac{1}{N}\sum_{i=1}^{N}(w_1 y_i \log p_i + w_0(1 - y_i)\log(1 - p_i)), \tag{5}$$

#### 6 Differential Evolution

In evolutionary computation, differential evolution (DE) [4, 5] is a method that optimizes a problem by iteratively trying to improve a candidate solution with regard to a given measure of quality.

DE is used for multidimensional real-valued functions but does not use the gradient of the problem being optimized, which means DE does not require the optimization problem to be differentiable, as is required by classic optimization methods such as gradient descent and quasi-newton methods. DE can therefore also be used on optimization problems that are not even continuous, are noisy, change over time, etc [3].

DE optimizes a problem by maintaining a population of candidate solutions and creating new candidate solutions by combining existing ones according to its simple formulae, and then keeping whichever candidate solution has the best score or fitness on the optimization problem at hand. In this way, the optimization problem is treated as a black box that merely provides a measure of quality given a candidate solution and the gradient is therefore not needed.

The steps of DE are described in Algorithm 2.

**Algorithm 2** Differential Evolution

$\alpha \leftarrow$ mutation rate
$popsize \leftarrow$ desired population size
$P \leftarrow \{\}$
$Q \leftarrow \{\}$
**for** $popsize$ times **do**
    $P \leftarrow P \cup \{new\ random\ individual\}$
**end for**
$Best \leftarrow null$
**while** Best is the ideal solution or time is over **do**
    **for** each individual $P_i \ \epsilon \ P$ **do**
        CalculateFitness($P_i$)
        **if** $Q \neq \{\}$ and $Fitness(Q_i) > Fitness(P_i)$ **then**
            $P_i \leftarrow Q_i$
        **end if**
        **if** $Best = null$ or $Fitness(P_i) > Fitness(Best)$ **then**
            $Best \leftarrow P_i$
        **end if**
    **end for**
    $Q \leftarrow P$
    **for** each individual $Q_i \ \epsilon \ Q$ **do**
        $\overrightarrow{a} \leftarrow$ a copy of individual other than $Q_i$, chosen at random with replacement from $Q$
        $\overrightarrow{b} \leftarrow$ a copy of individual other than $Q_i$ or $\overrightarrow{a}$, chosen at random with replacement from $Q$
        $\overrightarrow{c} \leftarrow$ a copy of individual other than $Q_i$ or $\overrightarrow{a}$ or $\overrightarrow{b}$, chosen at random with replacement from $Q$
        $\overrightarrow{d} \leftarrow \overrightarrow{a} + \alpha(\overrightarrow{b} - \overrightarrow{c})$
        $P_i \leftarrow$ one child from Crossover($d$,Copy($Q_i$))
    **end for**
**end while**
return Best

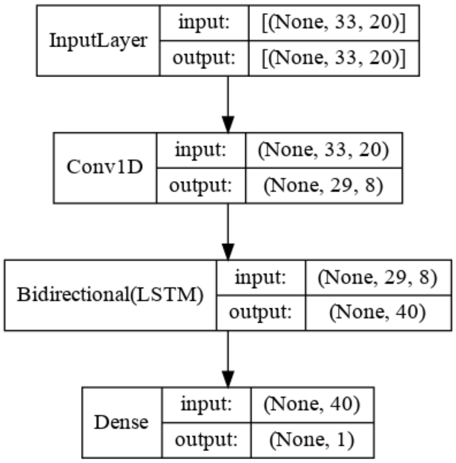


Figure 2: The one-hot encoded protein sequences are first fed into 1D convolution layer. The local features extracted from the CNN layer are then fed into the Bidirectional LSTM layer. The output of the bidirectional LSTM layer is fed into a fully connected dense layer with 1 neuron which will output the probability of the centered lysine residue of the input protein sequence to be succinylated.

### 7 Deep Learning Architectures

#### 7.1 CBL

The architecture of *BLC* is shown in Figure 2. The input layer takes as input the one-hot encoded protein sequences. 1D-CNN layer has been used with $kernel\ size = 5$, $strides = 1$ and $filters = 8$. Hence, $33 \times 20$ dimensional vector is converted into a 29-dimensional vector ($33 - 5 + 1 = 29$) for each of the filters resulting in a $29 \times 8$ dimensional vector. Then, the bidirectional LSTM layer is applied with $units = 20$ resulting in a 40-dimensional (20 for forward direction, 20 for backward direction) vector. Finally, a 1 neuron dense layer with sigmoid activation function is applied which outputs a value between 0 and 1 representing the probability of the lysine residue to be succinylated.

#### 7.2 BLC

The architecture of *BLC* is shown in Figure 3. After the input layer, we use the bidirectional LSTM layer with *return_sequences* set to *True* so that each LSTM cell outputs the hidden states for each timestep. Therefore, the bidirectional

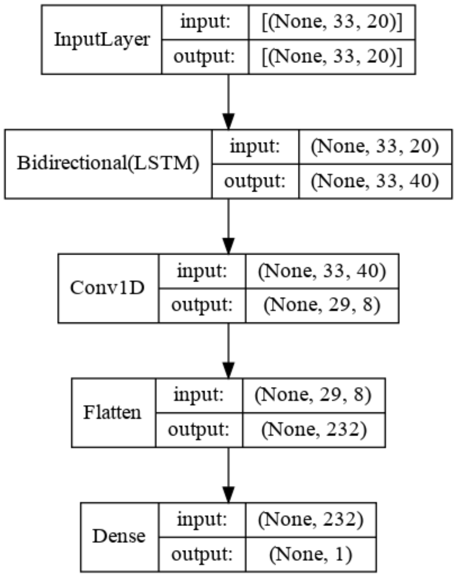


Figure 3: The one-hot encoded protein sequences are first fed into bidirectional LSTM layer. 1D-CNN layer follows this layer. The output of the CNN layer goes through a flatten layer. Finally, a fully connected dense layer outputs the probability in the same way as *CBL*.

LSTM layer outputs a $33 \times 40$ (there are 33 amino acids in a sample each of which represents a timestep for every LSTM unit) dimensional vector. Similar to the *CBL*, 1D-CNN layer outputs a $29 \times 8$ dimensional vector. As we will be feeding this vector into the feed forward network, we flatten the vector and feed it into the fully connected layer with 1 neuron as is done in *CBL*.

#### 7.3 CBL_BLC

The architecture of *CBL_BLC* is shown in Figure 4. There are two branches in this architecture. One branch is similar to *CBL* except for the fully connected layer, another branch is similar to *BLC* except for the fully connected layer. The outputs of the two branches are concatenated and fed into a fully connected network with two layers. The final layer has 1 neuron similar to the final layer of *CBL* and *BLC*.

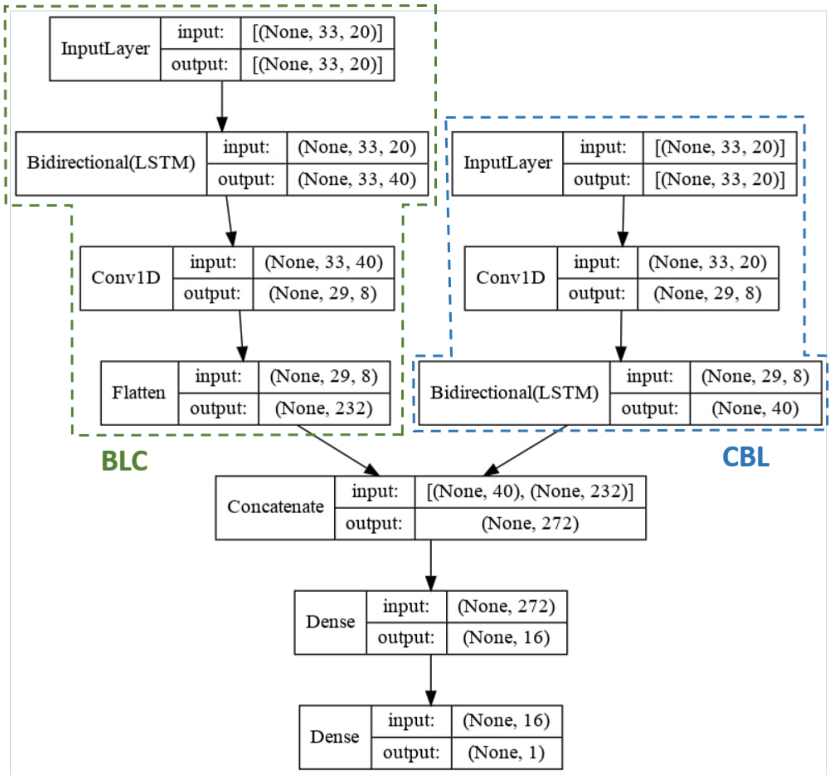


Figure 4: The *CBL_BLC* architecture has two branches. The left branch corresponds to the architecture of *BLC*. The one-hot encoded (details of one-hot encoding is discussed in Section 4 of Supplementary File) protein sequences are first fed into bidirectional LSTM layer. 1D-CNN layer follows this layer. The output of the CNN layer goes through a flatten layer. The right branch corresponds to the architecture of *CBL*. Here, the one-hot encoded protein sequences are first fed into 1D convolution layer. The local features extracted from the CNN layer are then fed into the Bidirectional LSTM layer. In CBL_BLC, the features from these two architectures are concatenated and passed through two densely connected layers.
